# Principles of Cell Circuits for Tissue Repair and Fibrosis

**DOI:** 10.1101/710012

**Authors:** Miri Adler, Avi Mayo, Xu Zhou, Ruth Franklin, Matthew Meizlish, Ruslan Medzhitov, Stefan Kallenberger, Uri Alon

## Abstract

Tissue-repair is a protective response after injury, but repetitive or prolonged injury can lead to fibrosis, a pathological state of excessive scarring. To pinpoint the dynamic mechanisms underlying fibrosis, it is important to understand the principles of the cell circuits that carry out tissue-repair. In this study, we establish a cell-circuit framework for the myofibroblast-macrophage circuit in wound-healing, including the accumulation of scar-forming extracellular matrix. We find that fibrosis results from multistability between three outcomes, which we term ‘hot fibrosis’ characterized by many macrophages, ‘cold fibrosis’ lacking macrophages, and normal wound-healing. The cell-circuit framework clarifies several unexplained phenomena including the paradoxical effect of macrophage depletion, the limited time-window in which removing inflammation leads to healing, the effects of cellular senescence, and why scar maturation takes months. We define key parameters that control the transition from healing to fibrosis, which may serve as potential targets for therapeutic reduction of fibrosis.

## Introduction

Tissue injury initiates a dynamic process that involves immune response, local proliferation of cells, scar deposition, and tissue regeneration. Following an injury, signals from the damaged cells of the tissue recruit inflammatory cells, such as monocytes and neutrophils, and stimulate differentiation of fibroblasts into myofibroblasts. Myofibroblasts produce extracellular matrix (ECM) proteins that form the scar, including the collagen and fibronectin. Recruited monocytes, together with tissue-resident macrophages, differentiate into inflammatory macrophages that break down and engulf the fibrin clot and cellular debris (Fig 1A). In a proper healing process, the inflammation is resolved within days, myofibroblasts and inflammatory macrophages are removed, the scar gradually degrades over weeks, and normal tissue composition is restored (Diegelmann and Evans, 2004; Duffield et al., 2013; Gurtner et al., 2008; Mann et al., 2011; Mutsaers et al., 1997).

**Figure 1:**
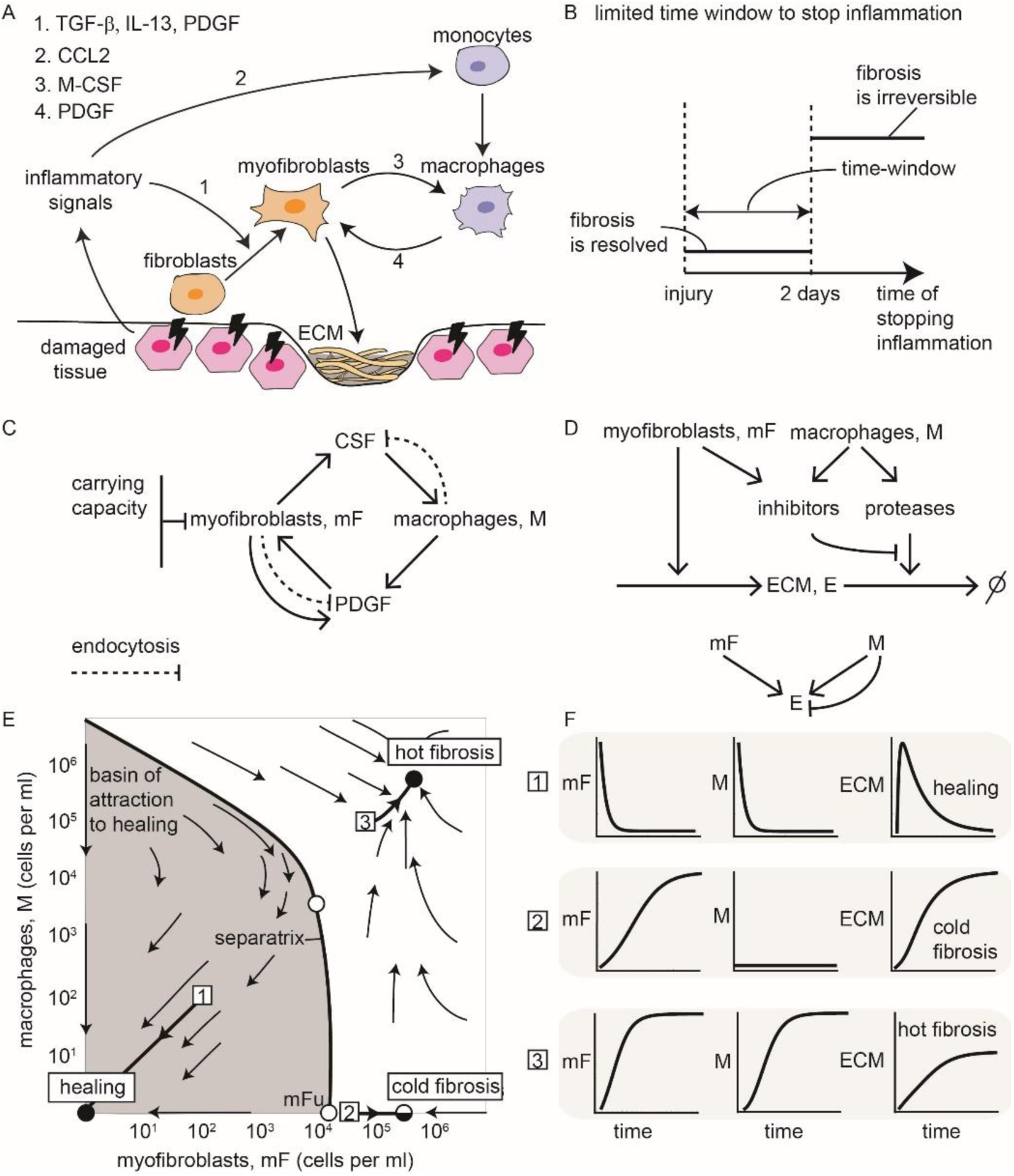
A circuit of communicating myofibroblasts and inflammatory macrophages shows multi-stability between a healing state and two fibrosis states. (A) Schematic process of wound healing and scar formation. (B) If injury is transient, it leads to brief inflammation, and no fibrosis occurs. However if injury is persistent and inflammation exceeds a critical time window, fibrosis is usually inevitable. (C) Circuit in which mF and M secrete growth factors (GFs) for each other. MFs show an autocrine loop and are limited by a carrying capacity. Both cell types remove the GFs by endocytosis (dashed arrows). (D) ECM is produced by mF and degraded by proteases mainly secreted by M. Proteases are inhibited by factors secreted by mF and M. (E) Phase portrait of the circuit, in which arrows show the flow rate of change of cell numbers. Stable fixed points (black dots) and unstable fixed points (white dots) are shown (the semi-stable cold fibrosis state as split black and white), as is the separatrix (black line) that marks the boundary between the basin of attraction of the healing (gray region) and fibrosis states. (F) Three temporal trajectories starting from the initial points 1-3 (white squares) are shown as well as the ECM accumulation.

However, protective tissue repair can evolve into fibrosis, a state of continuous scarring. Fibrotic scars begin to form within days and mature over months into an aggregate of ECM, myofibroblasts and macrophages. Fibrosis serves physiological purposes as a response to persistent insults. Excessive fibrosis can lead to fibrotic disease states, a common cause of age-related decline in organ function and ultimately of organ failure. Fibrotic diseases appear in many tissues including the skin, the musculoskeletal system, lung, liver, heart, brain, intestine and kidney (Wynn, 2008).

In many settings, proper repair occurs when the injury is transient and inflammation resolves quickly, whereas fibrosis appears if the injury is severe, prolonged or repetitive (Cao et al., 2016; Degryse et al., 2010; Mouratis and Aidinis, 2011). Studies show that there is a limited time-window of several days in which stopping inflammation avoids fibrosis (Fig 1B). Beyond this time window, fibrosis inevitably occurs even if inflammation is stopped. For example, in viral infection, early removal of the inflammatory trigger by antiviral drugs improves the patient state and reduces fibrosis (Ferguson and O’Kane, 2004; Wynn and Ramalingam, 2012). Similarly, early macrophage depletion substantially reduces the development of liver fibrosis (Duffield et al., 2005; Ide et al., 2005; Pradere et al., 2013; Sunami et al., 2012). The origin of this time window is unclear. In addition, the long timescales of scar maturation, which can take months, are surprising given this limited time-window, as well as the fact that cell turnover times are on the order of days.

In order to understand how a single process can lead to two very different physiological outcomes, healing and fibrosis, and to understand the timescales of fibrosis, it is important to define the circuit of cell-cell interactions that generates scarring. This circuit involves damaged tissue cells, myofibroblasts and macrophages. Damaged cells activate myofibroblasts, which produce ECM, and recruit monocytes which differentiate into macrophages. Monocyte-derived macrophages further activate myofibroblasts, and are important for the maintenance and resolution of fibrosis (Lech and Anders, 2013; Pakshir and Hinz, 2018; Wynn and Barron, 2010; Wynn and Vannella, 2016). These macrophages can transition between different states, including pro-inflammatory M1 and anti-inflammatory M2 states (Braga et al., 2015).

Macrophages and myofibroblasts reciprocally interact by growth factor exchange in which myofibroblasts secrete colony stimulating factor (CSF) for macrophages (CSF1, M-CSF) and macrophages secrete platelet-derived growth factor (PDGF) for myofibroblasts (Davies et al., 2013; Joshi et al., 2019; Lodyga et al., 2019; Wynn and Barron, 2010) (Fig 1A). In addition, myofibroblasts secrete PDGF in an autocrine loop which allows them to survive and expand in the absence of other growth-factor sources (Bonner, 2004; Trojanowska, 2008) (Fig 1C). This growth-factor communication is akin to the interaction between tissue-resident macrophages and fibroblasts that provides proper cell composition in the tissue, a circuit whose principles have been recently analyzed (Adler et al., 2018; Zhou et al., 2018).

Inflammatory macrophages also maintain the turnover of ECM by secreting factors that enhance or inhibit ECM degradation, namely matrix proteases (such as MMPs) and inhibitors of proteases (such as TIMPs). The latter are also secreted by myofibroblasts (Pellicoro et al., 2014) (Fig 1D). The secretion of antagonistic factors by macrophages may relate to the paradoxical effect of removing macrophages on fibrosis: depletion of macrophages can abrogate fibrosis or enhance it depending on context (Duffield et al., 2005).

Here, we present a cell-circuit framework for tissue repair and fibrosis. This framework clarifies how the interactions between the relevant cell types provide multi-stability: the dynamics can flow to the different physiological states of fibrosis or healing, depending on the severity and duration of the immune response. This approach builds on previous theoretical work regarding fibroblast activation (Hao et al., 2014), and adds to it the critical component of the fibroblast interaction with macrophages. The circuit predicts the existence of three steady-states: a state of healing associated with modest ECM production, and two fibrosis states associated with high cellularity and excessive ECM production, consistent with histopathological observations. In one of the fibrotic states, which we term ‘hot fibrosis’, myofibroblasts and macrophages are present at high levels. In the other state, which we term ‘cold fibrosis’, only myofibroblasts are present. The reciprocal macrophage-myofibroblast interaction is a key component in understanding observations such as the fibrotic time-window, the long timescale of scar maturation, and the paradoxical effect of macrophage depletion. Furthermore, the model suggests several targets for therapeutic reduction of fibrosis, including the myofibroblast autocrine loop.

## Results

### Macrophages and myofibroblasts interact to form a multi-stable circuit

We developed a circuit framework for the cell-cell interactions in wound healing (Fig 1A). Wound healing involves at least three main cell types. The first cell type consists of the tissue parenchymal cells, such as epithelial cells, that are damaged, and may eventually regenerate. These cells provide signals that recruit and activate the two other cell types: macrophages derived from circulating monocytes, and myofibroblasts derived from fibroblasts.

The presence of three cell types requires a three-cell circuit which describes the interactions among all three cell types. We find that a considerable simplification occurs if we focus on only two of the cell types, the macrophage-myofibroblast interactions, and consider the tissue parenchymal cells as a source of inflammatory signal that lasts as long as damaged cells are present. We thus consider the damage signal as an input that we can vary to explore transient versus repeated injury. The resulting two-cell circuit approach allows a clear understanding of the dynamics. In the SI we analyze the full three-cell circuit framework, and find that it leads to the same essential conclusions (Fig S1).

The two cell types, myofibroblasts and activated wound macrophages, communicate by secreting and sensing growth factors that are essential for their survival and promote proliferation. Myofibroblasts secrete CSF for the macrophages. Macrophages secrete PDGF for the myofibroblasts, as do the myofibroblasts themselves in an autocrine loop (Bonner, 2004). Here, we focus on monocyte-derived macrophages, and include their different states into a single variable. The growth factors are primarily removed via endocytosis by their target cells (Zhou et al., 2018).

We assume that myofibroblasts are close to carrying capacity – the maximum population size that can be supported in the tissue. In contrast, macrophages are far from their carrying capacity, as evidenced by the sharp rise in immune cell numbers during inflammation. A similar situation was found in a study of tissue homeostasis (Zhou et al., 2018).

We use this circuit to mathematically model the dynamics at the site of an injury. The variables are the numbers of myofibroblasts and macrophages at the injury site. The model takes into account secretion and consumption of growth factors as well as proliferation and removal of cells (Methods Eqs. 1-4). We used biologically plausible rate constants for secretion, endocytosis, proliferation and apoptosis based on in-vitro measurements (Zhou et al., 2018). Their values are listed in Table 1.

**Table 1:**
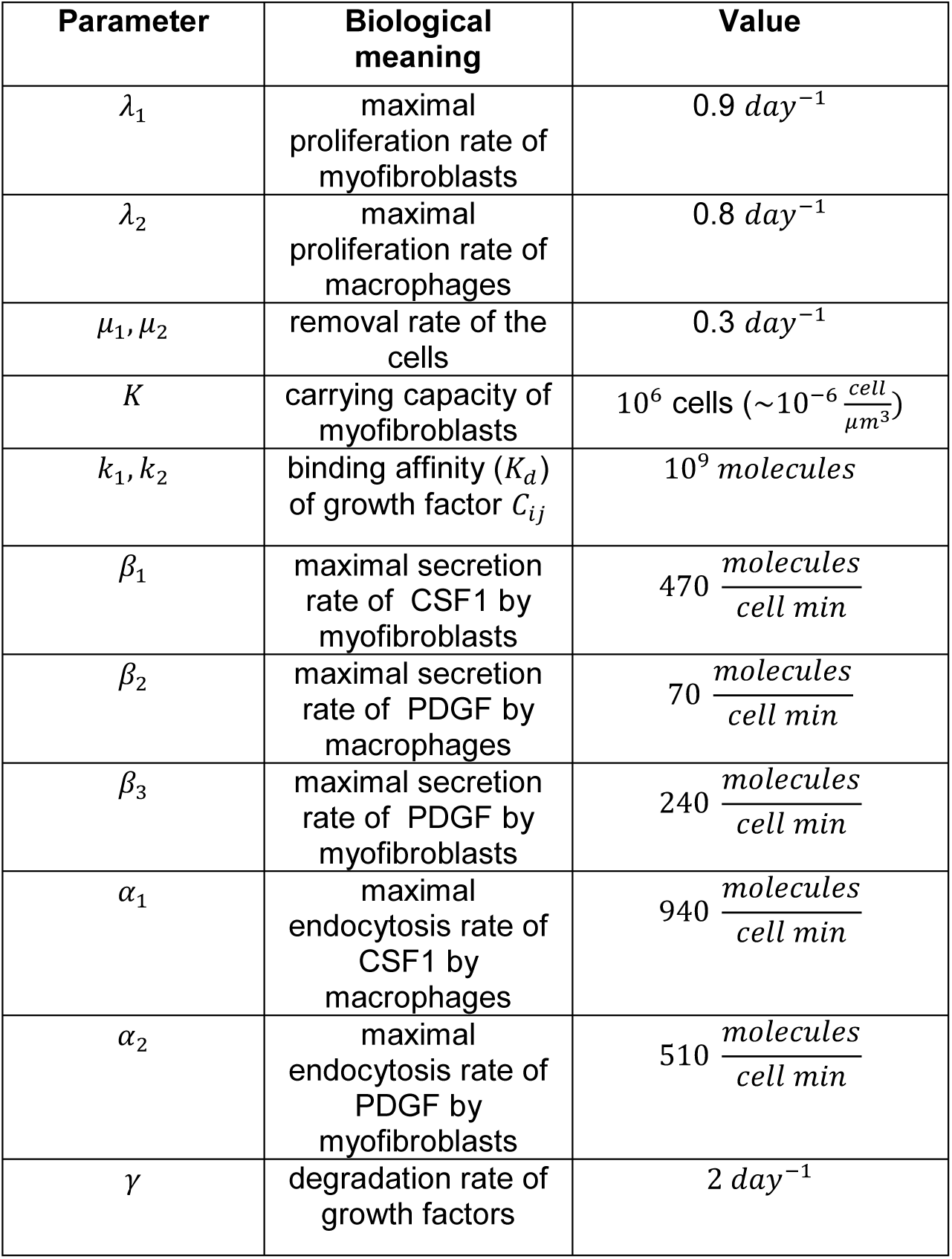
Plausible model parameter values.

| Parameter | Biological meaning | Value |
| --- | --- | --- |
| $\lambda_1$ | maximal proliferation rate of myofibroblasts | $0.9 \text{ day}^{-1}$ |
| $\lambda_2$ | maximal proliferation rate of macrophages | $0.8 \text{ day}^{-1}$ |
| $\mu_1, \mu_2$ | removal rate of the cells | $0.3 \text{ day}^{-1}$ |
| $K$ | carrying capacity of myofibroblasts | $10^6 \text{ cells } (\sim 10^{-6} \frac{\text{cell}}{\mu\text{m}^3})$ |
| $k_1, k_2$ | binding affinity ($K_d$) of growth factor $C_{ij}$ | $10^9 \text{ molecules}$ |
| $\beta_1$ | maximal secretion rate of CSF1 by myofibroblasts | $470 \frac{\text{molecules}}{\text{cell min}}$ |
| $\beta_2$ | maximal secretion rate of PDGF by macrophages | $70 \frac{\text{molecules}}{\text{cell min}}$ |
| $\beta_3$ | maximal secretion rate of PDGF by myofibroblasts | $240 \frac{\text{molecules}}{\text{cell min}}$ |
| $\alpha_1$ | maximal endocytosis rate of CSF1 by macrophages | $940 \frac{\text{molecules}}{\text{cell min}}$ |
| $\alpha_2$ | maximal endocytosis rate of PDGF by myofibroblasts | $510 \frac{\text{molecules}}{\text{cell min}}$ |
| $\gamma$ | degradation rate of growth factors | $2 \text{ day}^{-1}$ |

The resulting dynamics can be displayed in a phase plot, in which the axes are the concentrations of the two cell types (cells per ml of tissue volume). The arrows show the direction of change of the cell concentrations (Fig 1E).

The model shows three types of dynamic processes. At low initial cell (myofibroblasts + macrophages) concentrations, the dynamics flows towards a stable state with zero cells (Fig 1E). This state can be considered as the healing state, because the scarring process is resolved when monocyte-derived-macrophages and myofibroblasts are removed from the tissue, allowing the tissue-resident cell types to proliferate and restore tissue homeostasis.

However, if the initial cell numbers are high enough, above a threshold denoted by the separatrix line in Fig 1E, the dynamics flows to a steady state with high levels of both cell types. This steady-state is maintained with constant turnover of the cells. Myofibroblasts and macrophages keep each other at a high concentration due to the mutual secretion of growth factors. We consider this stable state as a ‘hot fibrosis’ state because both myofibroblasts and macrophages linger within the tissue and there is constant production of ECM. The term ‘hot’ indicates the high abundance of immune cells (macrophages).

The model also has a third fixed point. In this fixed point, there are zero macrophages and a high level of myofibroblasts. This state leads to ECM production because of the high level of myofibroblasts, and may be considered as another state of fibrosis. We term this ‘cold fibrosis’ due to the lack of macrophages.

The hot fibrosis and the healing fixed points are stable for a wide range of model parameters (Fig S2). The cold fibrosis state is robustly stable to changes in myofibroblast numbers but unstable regarding changes in the abundance of macrophages. A small influx of macrophages causes propagation to the hot fibrosis state. The cold fibrosis state can be stable in several realistic scenarios in which additional interactions are present or parameters are altered, as discussed in the SI (Fig S3).

To chart the amount of ECM during these dynamics, we computed the accumulation of ECM produced by myofibroblasts. We took into account ECM degradation by proteases including MMPs, and the inhibition of proteases, by factors including TIMPs (Fig 1D). Myofibroblasts enhance the accumulation of ECM by both producing ECM and inhibiting its degradation. Macrophages, in contrast, have a paradoxical effect on ECM degradation, by both degrading it and inhibiting its degradation (Fig 1D) (Methods Eqs. 5-11). The resulting amount of ECM is shown in Fig 1F. The healing state has a small amount of ECM accumulation followed by degradation, whereas the hot fibrosis and the cold fibrosis states have high persistent level of ECM (Fig 1F).

Although we model ECM concentration, the same variable may be interpreted to include also matrix quality such as stiffness (Klingberg et al., 2013). Often, excess matrix is stiffer, abnormal and partially degraded. Such stiffness may accelerate myofibroblast differentiation and accumulation, as well as enhance the transition to fibrosis (Avery et al., 2018).

### Bistability of the circuit explains how persistent injury triggers fibrosis

To study the effect of the type of injury on fibrosis, we considered the response of the myofibroblast-macrophage circuit to several scenarios of injury. We studied injury stimuli consisting of a single transient pulse, repetitive injury pulses, and a single prolonged pulse. We modelled these injury stimuli by inflammatory signals that recruit monocyte-derived-macrophages, and thus serve as a source term in the macrophage equation (Methods Eq. 12).

We solve the dynamical trajectory of the cells in response to these injuries starting from an initial condition of a small number of macrophages and myofibroblasts (1 cell/ml). The results do not depend sensitively on this initial condition. We find that the outcome of the cell-circuit response depends on the severity, duration and recurrence of the injury.

A transient injury stimulates a brief inflammatory pulse (Fig 2A) that recruits monocytes to differentiate into macrophages. We modeled this as a strong, brief influx of macrophages. When the pulse ends, the macrophages die, and no longer support myofibroblast proliferation. Both cell types decline to zero and reach the healing stable state (Fig 2B). The overall accumulation of ECM is very small (Fig 2G).

**Figure 2:**
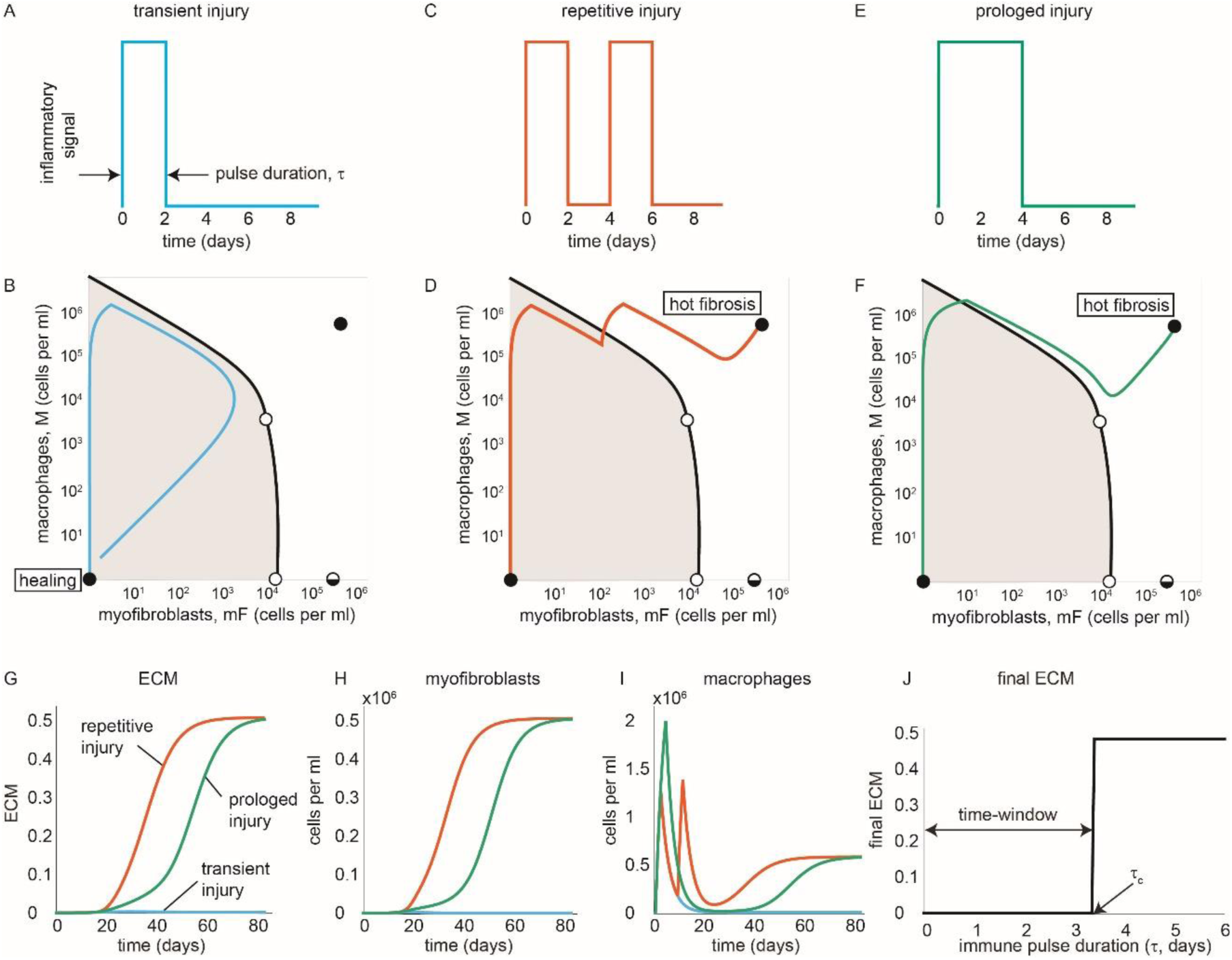
The mF-M circuit shows healing versus fibrosis depending on duration and recurrence of inflammation. (A) For a brief pulse of inflammation (2 days), the rise of mF and M is transient, leading to a healing trajectory in phase space that returns to the healing state (in light blue) (B). In contrast, two successive 2-days long inflammatory pulses (C), or a prolonged pulse (4 days) (E) lead to a trajectory to the hot fibrosis state with persistent mF and M populations (in red and green, respectively) (D, F). Note that the separatrix applies to the equations without the external inflammation input, and so the separatrix can be crossed during the input pulse(s). The dynamics of ECM (G), mFs (H) and Ms (I) are plotted in response to transient (light blue lines), repetitive (red lines), and prolonged (green lines) injuries. (J) Final ECM as a function of inflammation pulse duration shows a critical time-window of about 3 days to stop inflammation.

In contrast, a repetitive (Fig 2C), or prolonged injury (Fig 2E) leads to a different result. The cells divide and grow, mutually supporting each other via growth factors, until they cross the separatrix between the healing and fibrosis states. The cells converge to the hot fibrosis state (Fig 2D, F). Here, ECM accumulation is enhanced and the ECM reaches a high steady-state which represents the excess scar formation in fibrosis (Fig 2G).

Brief repetitive injuries have some dynamical differences from a single prolonged injury. When the prolonged injury elicits a response that is close to the separatrix, the scar shows a slower accumulation than reparative injury (Fig 2G, green line compared to red line) due to the temporary loss of macrophages (Fig 2I). Very prolonged injury becomes more similar to repetitive injury (Fig S4).

To illustrate the existence of a critical time-window, we plot the final ECM level as a function of the duration of a transient immune pulse. The final ECM level is very small up to a critical pulse duration of *τ*_*c*_∼3 days. For longer pulses, ECM jumps to a high final level, representing fibrosis (Fig 2J). The critical time-window is due to the crossing of the separatrix. Thus, the model predicts that inflammation must be stopped early in order to avoid fibrosis.

### The timescale of months for fibrotic scar maturation is due to the slow crossing of a dynamical barrier

We also analyzed the time to reach full ECM levels, which corresponds to the scar maturation time. We define the maturation time (*t*_*m*_) as the time when half of the final ECM level has been accumulated (Fig 3A). The maturation time is very long for immune pulses close to the critical pulse duration, and reaches 80 days (Fig 3B). For longer immune pulses, maturation time is about a month. This month-timescale is surprising, because it is much longer than the turnover time of the growth factors (hours) or the cells (days).

**Figure 3:**
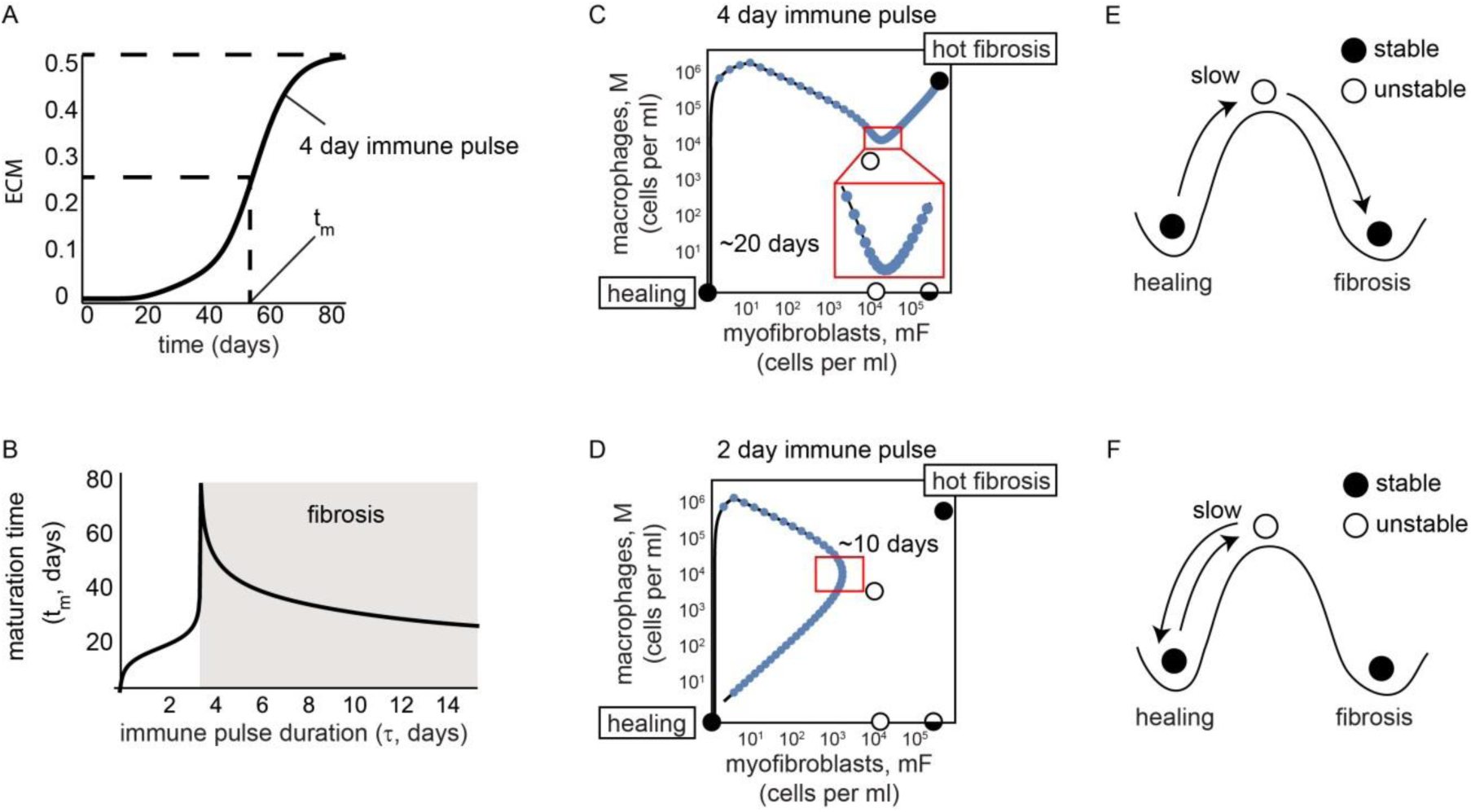
The mF-M circuit explains the timescale of months for scar maturation. (A) ECM accumulation in response to a 4 day immune pulse. Maturation time, *t*_*m*_, is defined as the time to reach half maximal ECM accumulation. (B) ECM maturation time is on the order of months for fibrosis, and weeks for healing. The slow timescale of weeks-months is due to a dynamical barrier due to an unstable fixed point (upper white circle) (C-D), akin to a ball slowing down at the top of a hill (E-F). In (C, D) blue dots indicate trajectory values at intervals of one day, so that slow dynamics correspond to dense blue dots.

To understand the slow maturation time, we zoomed in on the dynamics for an immune pulse of 4 days (Fig 3C). During the pulse, there is a sharp rise in macrophages due to influx from monocytes. Because the pulse is brief, there are only a few myofibroblasts when the pulse ends, and they cannot strongly support the macrophages by CSF secretion. Hence the macrophage population drops after the pulse is over, but does not vanish.

The dynamics are then close to an unstable fixed point (upper white circle in Fig 3C). Exactly at the fixed point the rates of change are, by definition, zero. Therefore, close to the fixed point the rates of change are small, and the system slowly moves away from the fixed point by means of cell growth. Intuitively, the tissue crosses a dynamical threshold from the healing to the hot fibrosis state, akin to a ball that slows down when it reaches the peak of a hill (an unstable fixed point), from which it descends to a new fixed point (Fig 3E). Indeed, the dynamics creep up slowly towards the hot fibrosis state, but it takes about 20 days to appreciably move towards this mature final state (Fig 3C).

Interestingly, the slow timescale is also seen when healing occurs in the model. For an immune pulse of 2 days (Fig 3D), which is shorter than the critical window, the dynamics also come close to the unstable fixed point, but in this case the trajectory drops back to the ‘healing’ side of the hill (Fig 3F). Indeed, experiments show that non-fibrotic resolution of transient injury can take weeks (Seifert et al., 2012).

### Depletion of macrophages at different myofibroblast numbers show opposing effects

We next analyze the effect of macrophage depletion, which has been reported to be either pro- or anti-fibrotic in different experiments. Once macrophages are depleted, myofibroblast survival depends on their autocrine loop. This autocrine loop provides bistability in which myofibroblasts either converge to the cold fibrosis state and survive, or die and cause the tissue to flow to the healing state. The threshold for myofibroblasts, above which they can sustain their proliferation without macrophages, is the unstable fixed point mFu (lower white circle in Fig 4A).

**Figure 4:**
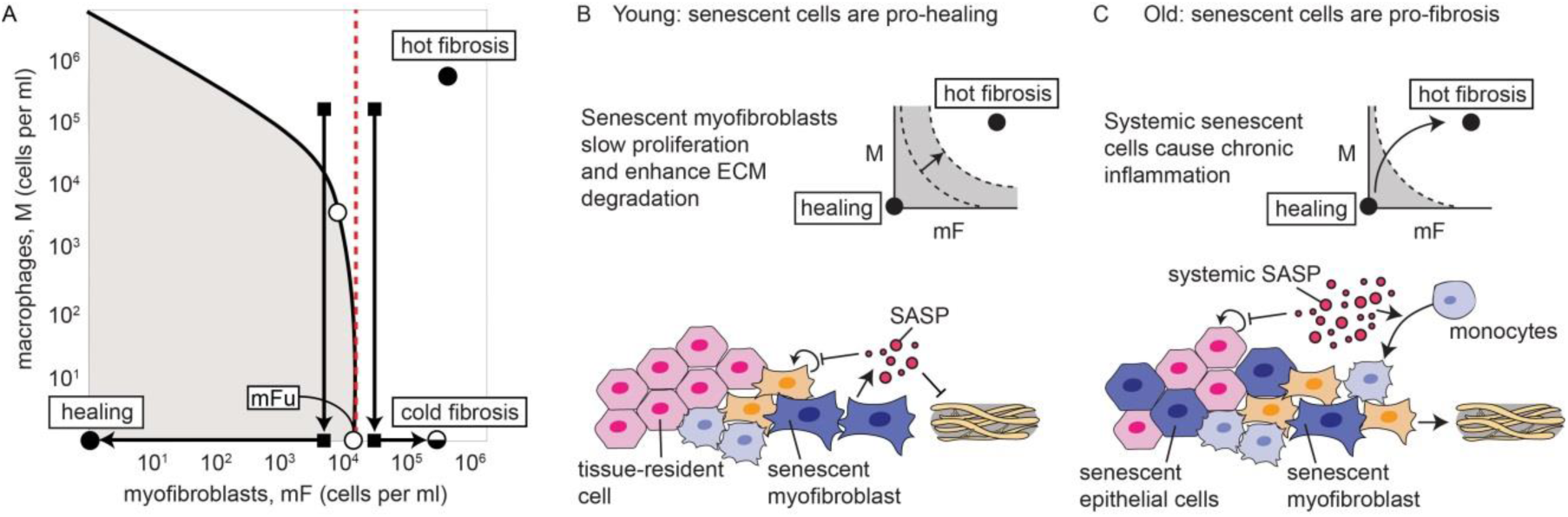
Paradoxical effects of macrophage depletion and senescent cells on fibrosis. (A) Depleting macrophages when myofibroblasts are below or above the mFu unstable point (lower white circle) leads to healing or fibrosis, respectively. (B) Role of SnCs is pro-healing in young individuals in which SnCs are mainly locally generated myofibroblasts at the injury site that enlarge the basin of attraction to the healing state. (C) At old ages, systemic SnCs cause chronic inflammatory signals which can cause the tissue to cross the separatrix.

This unstable fixed point can explain the paradoxical effects of removing macrophages on fibrosis. If macrophages are removed early enough, before myofibroblasts cross mFu, the myofibroblasts cannot support themselves, and the system flows to the healing state. In contrast, removing macrophages when myofibroblasts exceed mFu leads to a rapid flow to the cold fibrosis state characterized by high ECM (Fig 4A). This bistability may explain why removal of macrophages enhanced healing in some studies, whereas in other studies, removal led to accelerated fibrosis (Duffield et al., 2005), even though removal started at similar levels of macrophages.

### The model suggests different roles for senescent cells in young and old organisms

We can use the present approach to explore the impact of cellular senescence on tissue repair and fibrosis. Senescent cells (SnCs) are known to be pro-healing at young ages; for example, removing senescent cells impairs repair in the skin and liver (Jun and Lau, 2010; Krizhanovsky et al., 2008). On the other hand, senescent cells are pro-fibrosis at old ages and in age-related diseases. For example, removing whole-body SnCs helps reduce fibrosis in IPF models and in systemic sclerosis (Hecker et al., 2014; Muñoz-Espín and Serrano, 2014; Piera-Velazquez and Jimenez, 2015). The present model can offer a framework to understand this age-dependent role.

SnCs are cells that stop dividing and secrete factors, known as senescent-associated secretion profiles (SASP) that includes inflammatory signals, ECM degradation factors and factors that inhibit proliferation of nearby cells (van Deursen, 2014). The number of SnCs increases dramatically with age in many tissues. At all ages, senescent fibroblasts are generated at the injury site during the process of normal healing.

One may propose the following picture: at young ages, systemic SnC levels are low. SnCs are generated locally after injury, mainly senescent myofibroblasts. They enhance healing because they i) slow down myofibroblast proliferation by a bystander effect through SASP, ii) increase myofibroblast loss due to myofibroblast senescence and iii) degrade ECM. All of these factors enlarge the basin of attraction of the healing state (Fig 4B).

In contrast, at old ages, SnCs are abundant in many tissues even without injury (Campisi, 2005; Muñoz-Espín and Serrano, 2014), including senescent epithelial cells. These SnCs have systemic effect through circulating SASP. SnCs thus cause chronic inflammation. These systemic inflammatory signals, together with increased local senescence, can effectively prolong the immune signal that occurs upon injury, pushing it into the fibrotic basin of attraction (Fig 4C). The threshold for fibrosis is thus increasingly lowered with age.

In most people this threshold is not normally crossed. In people with genetic and environmental conditions that cause increased damage to a given tissue, fibrosis can occur if the damage exceeds the age-related threshold. This can lead to organ-specific fibrotic diseases that are strongly age related, and occur only in a fraction of the population.

Thus, the local nature of senescence at young ages versus the systemic nature of SnCs at old ages may contribute to the opposing effects of SnCs on fibrosis.

### Weakening the PDGF autocrine loop or slowing myofibroblast proliferation may prevent or reverse fibrotic response

Finally, we asked what interventions might prevent fibrosis. For this purpose, we asked how the basin of attraction for the healing state can be enlarged. Such enlargement means that more situations will end up resolved without fibrosis.

We find that prevention occurs in the model when the cold fibrosis fixed-point vanishes. The parameters that control the existence of the cold fibrosis fixed point are PDGF autocrine secretion (*β*_3_) and endocytosis (*α*_2_) rates and the myofibroblast proliferation (*λ*_1_) and removal (*μ*_1_) rates. Increasing the PDGF autocrine endocytosis rate or myofibroblast removal rate, or decreasing PDGF autocrine secretion rate or myofibroblast proliferation rate, removes the cold fibrosis fixed point. This provides a larger time window in which removing inflammation leads to proper healing (Fig 5A-B). The requirement for such prevention of fibrosis (removal of the fixed point) can be summarized as a single requirement on a dimensionless combination of these four parameters: 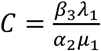 needs to be smaller than 1, as can be analytically shown(Methods). The cold fibrosis vanishes also for a larger PDGF degradation rate (*γ*) or a larger binding affinity of PDGF to its cognate receptor (*k*_2_) (Fig S2).

**Figure 5:**
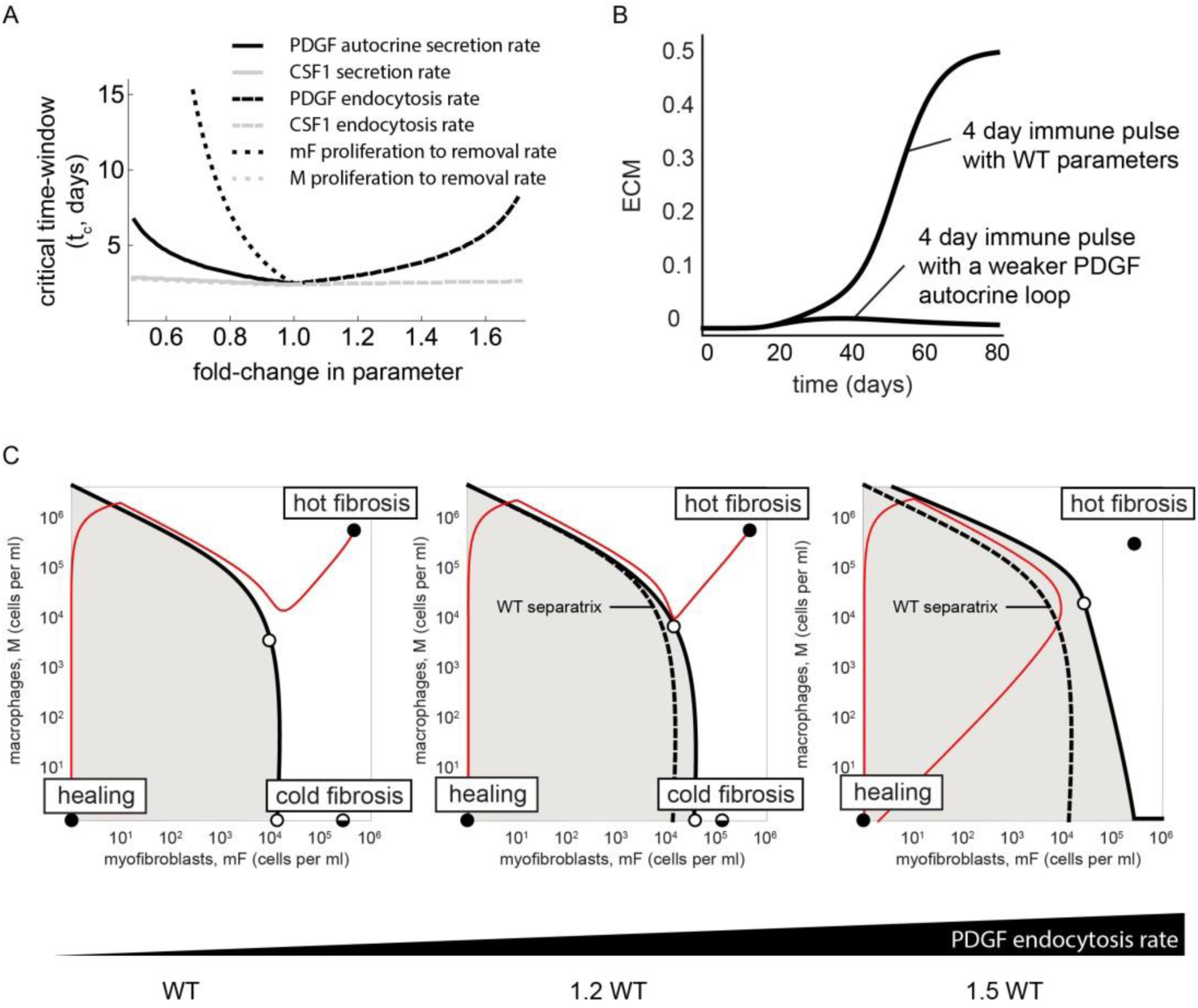
Fibrosis can be prevented or reversed by changing several circuit parameters. (A) Critical time window for inflammation that results in healing as a function of fold-change in circuit parameters. Only changes that lengthen the time window are shown. The window can be lengthened by decreasing autocrine secretion, increasing PDGF endocytosis, or decreasing the ratio of mF proliferation to removal. With altered parameters, even a 4-day immune pulse leads to healing, with low ECM production (B). (C) Increasing PDGF endocytosis rate by 150% eliminates the cold fibrosis state, enlarging the basin of attraction to the healing state (gray region).

The dynamics when this condition is met (*C* = 0.9 and thus *C* < 1) are shown in Fig 5B. A lengthy immune pulse of 4 days (Fig 5B), that would lead to fibrosis with wild-type parameters, now flows to the healing state with no fibrosis (Fig 5B). The hot fibrosis state can still be reached after a pulse of 6 days.

We also asked whether fibrosis can be reversed, in the sense that a mature scar in the hot fibrosis state can be made to flow to the healing state by depleting macrophages. This can be achieved if both myofibroblasts and macrophage are dropped below the separatrix. One approach is to first achieve the prevention criterion above, *C* < 1, eliminating the cold fibrosis state. This means that myofibroblasts can no longer persist in the absence of macrophages. If then macrophages are depleted enough to drop below the separatrix, the system must flow to the healing state no matter how many myofibroblasts are present (Fig 5C). Such macrophage depletion treatment can be short-term: once the separatrix is crossed, the circuit will flow to the healing state even if one stops depleting macrophages. The predicted reversal of fibrosis depends on the fact that fibrosis is a dynamic steady-state with cell turnover.

## Discussion

This study presented a circuit for wound healing and fibrosis based on the interactions between myofibroblasts and inflammatory macrophages. The circuit explains how two drastically different outcomes result from the same system: healing when injury is transient and fibrosis with excessive ECM production when the injury is prolonged or repetitive. Each of these outcomes has its own basin of attraction in a bistable phase diagram (Fig 6A). We use the cell-circuit framework to explain several physiological and pathological phenomena in tissue repair and fibrosis. First, we show that the cell-circuit response depends on the duration of the inflammatory signal, which explains the origin for a brief critical time-window in which removing inflammation abrogates fibrosis. Second, we show that the months-long timescale for scar maturation, which is surprising given the much shorter timescales of cell signaling and proliferation (hours-days), originates from a slowdown near an unstable fixed point. Thirdly, we explain the paradoxical effect of macrophage depletion: healing or fibrosis occur when myofibroblasts are below or above a critical threshold, respectively, where they can support their own proliferation at the time of macrophage depletion. Finally, we suggest potential targets for preventing and reversing fibrosis including the myofibroblast autocrine loop.

**Figure 6:**
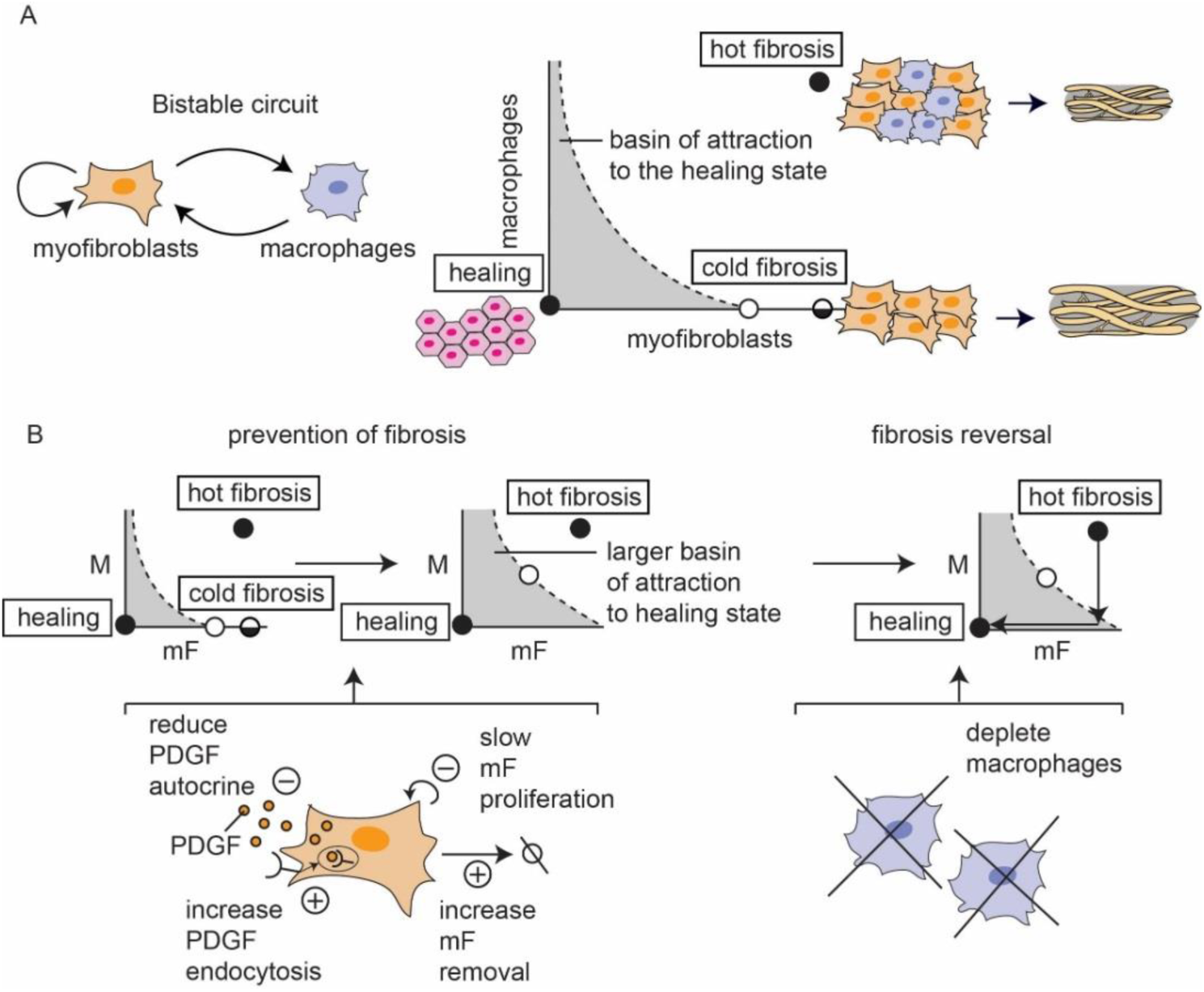
Overview of the present circuit approach to understanding healing and fibrosis. (A) The M-mF circuit shows a stable healing state and two fibrosis states. Outcome depends on the duration and persistence of inflammation pulses caused by an injury, which can remain in the basin of attraction for healing, or cross the separatrix into the basin of attraction for fibrosis. (B) The present analysis suggests targets for prevention and reversal of fibrosis, by eliminating the cold fibrosis fixed point and enlarging the basin of attraction to the healing state.

The present circuit framework indicates that constant cell turnover is needed to maintain fibrosis states. It explains why several types of fibrosis are sometimes seen in different clinical settings (Soderblom et al., 2013). The circuit shows two types of fibrosis, hot fibrosis with both macrophages and myofibroblasts, and cold fibrosis with only the latter (Fig 6A). The cold fibrosis fixed point explains why the depletion of macrophages can sometimes result in accelerated fibrosis (Duffield et al., 2005).

Parallels can be drawn to several pathologies resulting in fibrosis. In fibrotic diseases caused by persistent damaging agents, such as pneumoconiosis or liver fibrosis, fibrotic lesions typically show infiltrates of myofibroblasts as well as macrophages (Castranova and Vallyathan, 2000; Lech and Anders, 2013; Pellicoro et al., 2014; Tesch, 2010). This resembles hot fibrosis.

Analysis of the basins of attraction of a model with stable cold fibrosis suggests that very prolonged injury, such as damage signals generated by damaging agents that cannot be removed, tends to lead to hot fibrosis (Fig S3B). Injury signals of intermediate duration tend to lead to cold fibrosis (Fig S3B).

Hot and cold fibrosis may also correspond to the tissue composition of pathological scars in dermatology. The two main types of pathological scars in the skin are hypertrophic scars and keloid (Abergel et al., 1985; Cohen et al., 1972; Craig et al., 1975; Ogawa, 2017; Trace et al., 2016). Keloids are characterized by high densities of macrophages, as in hot fibrosis, and inflammation persists over years (Santucci et al., 2001). In contrast, the transition from early to late hypertrophic scars is accompanied by a progressive decrease of immune cell infiltrates, as in cold fibrosis (Santucci et al., 2001).

Keloids (hot fibrosis) can be treated by anti-proliferative therapies such as local injection of cortisol, cryotherapy, radiotherapy or topical application of cytostatic drugs (Arno et al., 2014). An exclusively surgical treatment of keloids results in regrowth (Love and Kundu, 2013), in contrast to hypertrophic scars. Based on the model, moving from hot fibrosis to cold fibrosis by a cytostatic treatment that affects activated macrophages brings the system to a state closer to hypertrophic scars; this can be followed by surgery. This conclusion is coherent with the observation that combining keloid surgery with anti-proliferative treatment decreases recurrence (Arno et al., 2014).

Extensions to the present model can include different macrophage states (such as M1 and M2) and additional growth factor interactions. Even with these extensions, it is likely that the core mechanism of bistability will allow the circuit to show healing versus fibrosis. It would be interesting to explore how this circuit might interface with cancer, to model the fibrotic-like microenvironment that promotes cancer incidence and cancer growth (Dvorak, 1986; Kalluri, 2016; Marsh et al., 2013).

The present circuit provides possible therapeutic targets for fibrosis, such as the PDGF autocrine loop for myofibroblasts. Intriguingly, enhancing PDGF autocrine secretion rate is predicted to worsen fibrosis. Indeed, over-expressing PDGF*α* in mouse myofibroblasts increased the probability of fibrosis (Olson and Soriano, 2009). In the present model, fibrosis can be prevented by any of the following or their combination: reduction of PDGF autocrine secretion rate, reduction of myofibroblast proliferation rate, increase in PDGF endocytosis or degradation rate, increase in PDGF binding affinity or increase in myofibroblast removal rate. Fibrosis can also be reversed according to the present picture. To reverse fibrosis requires altering any of the above parameters, in order to abrogate the cold fibrosis fixed point, together with depletion of macrophages from the mature scar, to cause the system to flow to the healing state.

More generally, the present circuit approach can be extended to suggest principles for additional physiological and pathological processes, in order to define essential dynamical mechanisms and generate insights for experiments.

## Methods

### Model for the myofibroblast-macrophage circuit

We model the reciprocal interaction between monocyte-derived-macrophages and the ECM-producing myofibroblasts. We assume a mean-field approximation where all cells see the same concentration of growth factors. Macrophages (*M*) secrete PDGF for myofibroblast survival and division, and myofibroblasts (*mF*) secrete CSF for macrophage survival and division (Fig 1C). The balance of cell proliferation and removal yields equations for the rate of change of cell numbers:

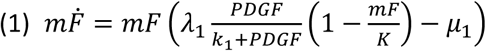

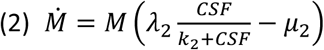

where *λ*_1_, *μ*_1_, *λ*_2_, *μ*_2_ are the proliferation (at saturating GF) and removal rates of myofibroblasts and macrophages, respectively. The effect of each growth factor on its target cell occurs by binding of the growth factor to its cognate receptor on the target cells, as described by Michaelis–Menten functions with halfway effect at *k*_1_ for PDGF receptor and *k*_2_ for CSF receptor. Myofibroblasts have a carrying capacity, *K*– the maximum cell population size that can be supported in the tissue, whereas macrophages are assumed to be far from their carrying capacity.

The equations for the growth factors include production terms, removal by endocytosis and degradation terms, and include autocrine production of PDGF by myofibroblasts:

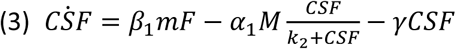

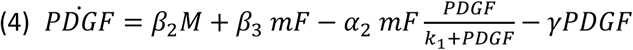

Here, CSF is produced by myofibroblasts at rate *β*_1_, and endocytosed by macrophages at maximal rate *α*_1_ (Eq. 3). PDGF is produced by macrophages and myofibroblasts at rates *β*_2_ and *β*_3_ respectively, and endocytosed by myofibroblasts at maximal rate *α*_2_ (Eq. 4). Both growth factors are degraded at rate *γ*. We assume here that endocytosis works with Michaelis–Menten kinetics with the same halfway points, *k*_*i*_, as the effect of the growth factors on their target cells in Eqs. 1 and 2, because both signaling and endocytosis depend on ligand binding to the cognate receptor. The parameter values that we used are listed in Table 1, using the measurements and estimates of ref (Zhou et al., 2018).

### Model ECM accumulation

We model the dynamical accumulation of ECM (E) which is produced by myofibroblasts. ECM degradation is controlled by proteases, P, such as MMPs, which are inhibited by anti-proteases A (such as TIMPs). P are secreted mainly by macrophages and A are produced by both macrophages and myofibroblasts, resulting in:

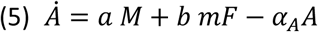

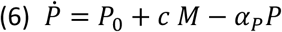

Here, A are produced by macrophages and myofibroblasts at rates a and b, respectively, and degraded at rate *α*_*A*_ (Eq. 5). We assume basal proteases secretion by other cell types, *P*_0_. P are produced mainly by macrophages at rate c and degraded at rate *α*_*P*_. Assuming that P and A reach their steady-states on a faster timescale than the cells and ECM, we calculate their steady-states by equating Eqs. 5-6 to zero:

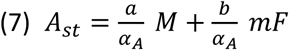

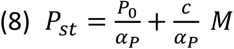

We next consider the equation for ECM, where P enhance ECM degradation and A inhibit it:

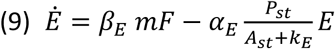

Here, *β*_*E*_ is the production rate of ECM by myofibroblasts and *α*_*E*_ is its maximal removal rate. *k*_*E*_ is the halfway point of inhibition of ECM degradation by A. Plugging in the P and A steady states (Eqs 7-8) in Eq. 9 yields:

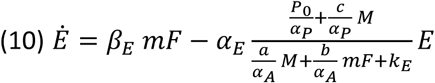

We use dimensional analysis in which we define dimensionless variables: 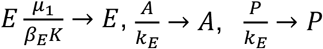 and dimensionless parameters: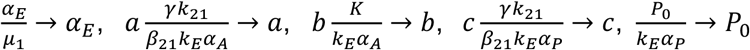. Using these dimensionless variables and parameters Eq. 10 now reads:

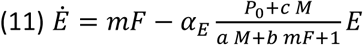

Thus, the steady state of ECM is: 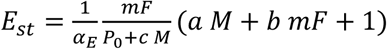. The parameter values that we used for the ECM dynamics are listed in Table 2.

**Table 2:** Dimensionless parameter values for ECM dynamics.

| Parameter | Definition | Value |
| --- | --- | --- |
| $a$ | $a \frac{\gamma k_{21}}{\beta_{21} k_E \alpha_A}$ | 1 |
| $b$ | $b \frac{K}{k_E \alpha_A}$ | 1 |
| $c$ | $c \frac{\gamma k_{21}}{\beta_{21} k_E \alpha_P}$ | 1 |
| $P_0$ | $\frac{P_0}{k_E \alpha_P}$ | 0.1 |
| $\alpha_E$ | $\frac{\alpha_E}{\mu_1}$ | 1 |

### Simulating inflammatory signals

We model different injury scenarios by considering a pulsatile inflammatory signal, *I*(*t*), that positively controls the production rate of macrophages. The equation for macrophages thus reads:

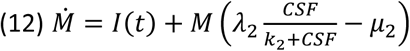

The pulsatile inflammatory signal is given by:

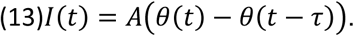

Therein, θ(*t*) denotes the Heaviside step function. We simulated three different signals. For the transient injury, we consider a single pulse with amplitude 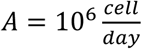 and duration of *τ* = 2 days. For the repetitive injury, we consider two successive pulses ofamplitude 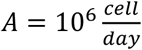 and duration of *τ* = 2 days. For the prolonged injury, we consider a single pulse with 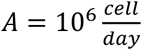 and duration of *τ* = 4 days.

### Parameters for fibrosis prevention and reversal by eliminating the cold fibrosis fixed point

To analyze the stability of the cold fibrosis fixed point, we consider the situation with zero macrophages. Therefore the equations that describe myofibroblast dynamics include myofibroblasts and their autocrine secreted growth factor PDGF. Near the cold fibrosis fixed point, PDGF is mainly removed by endocytosis by myofibroblasts and we can neglect its non-endocytotic degradation rate *γ*:

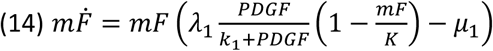

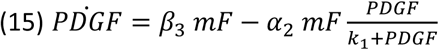

Using a quasi-steady state approximation due to the faster timescale of GFs compared to cells, we calculate the PDGF steady state:

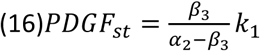

Substituting this in the equation for myofibroblasts we now have:

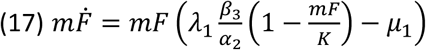

Solving for the steady state of myofibroblasts by equating Eq. 17 to zero yields either the healing state (*mF* = 0) or the approximate cold fibrosis fixed point:

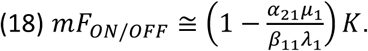

The expression (Eq. 18) provides a condition for the existence of the cold fibrosis fixed point. Since myofibroblast number cannot be negative, the ratio,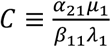 must besmaller than 1 in order to have a solution except for the zero solution. This means that in case *C* is large enough, the cold fibrosis state will not be a solution and the flow is to the healing fixed point.

