## Supplementary Information for "Principles of Cell Circuits for Tissue Repair and Fibrosis"

### Myofibroblasts, damaged epithelial cells, and inflammatory macrophages interact to form a multi-stable circuit

We model the interactions between damaged epithelial cells ($D$), myofibroblasts ($mF$), and macrophages ($M$) using the same equations for myofibroblasts and for the growth factors (Eqs. 1, 3-4) and the following equations for macrophages and the damaged epithelial cells:

(S1) $\dot{M} = D + M\left(\lambda_2 \frac{CSF}{k_2+CSF} - \mu_2\right)$

(S2) $\dot{D} = d(t)(N - D) - \alpha\, D$

where $d(t) = d_0\big(\theta(t) - \theta(t-\tau)\big)$ is the damage stimuli, $N$ is the total concentration of epithelial cells in the tissue including normal and damaged cells, and $\alpha$ is the removal rate of the damaged epithelial cells.

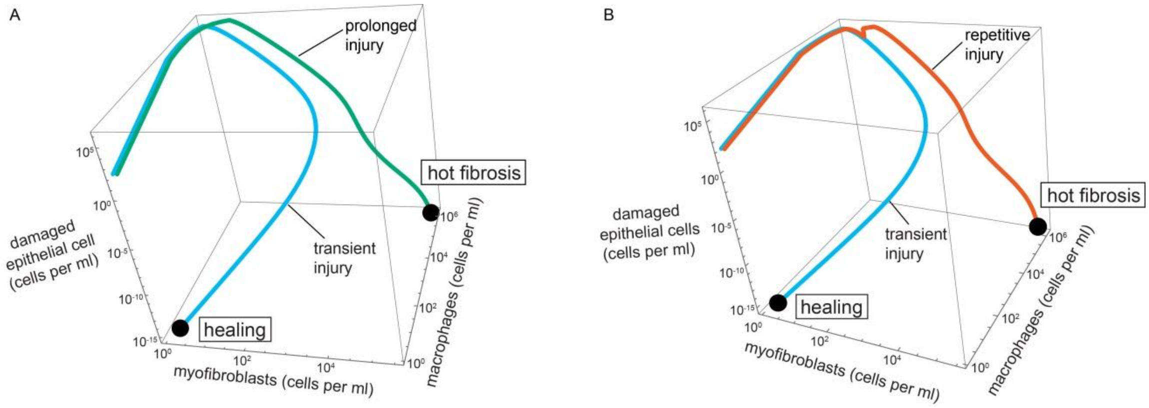


**Figure S1: A three-cell-type circuit of communicating myofibroblasts, damaged epithelial cells, and inflammatory macrophages shows healing versus fibrosis depending on duration and recurrence of injury.** (A) A transient injury that cause damage to epithelial cells leads to accumulation of myofibroblasts and macrophages followed by their removal and removal of damaged epithelial cells (light blue trajectory). In contrast, prolonged injury (green trajectory, A) or repetitive injury (red trajectory, B) causes the circuit dynamics to flow towards the hot fibrosis state with high levels of myofibroblasts and macrophages with excess ECM, after the damaged epithelium is removed. We used the parameter values: $N = 10^6\ cells$, $\alpha = 1\frac{1}{day}$, $d_0 = 100\frac{cells}{day}$.

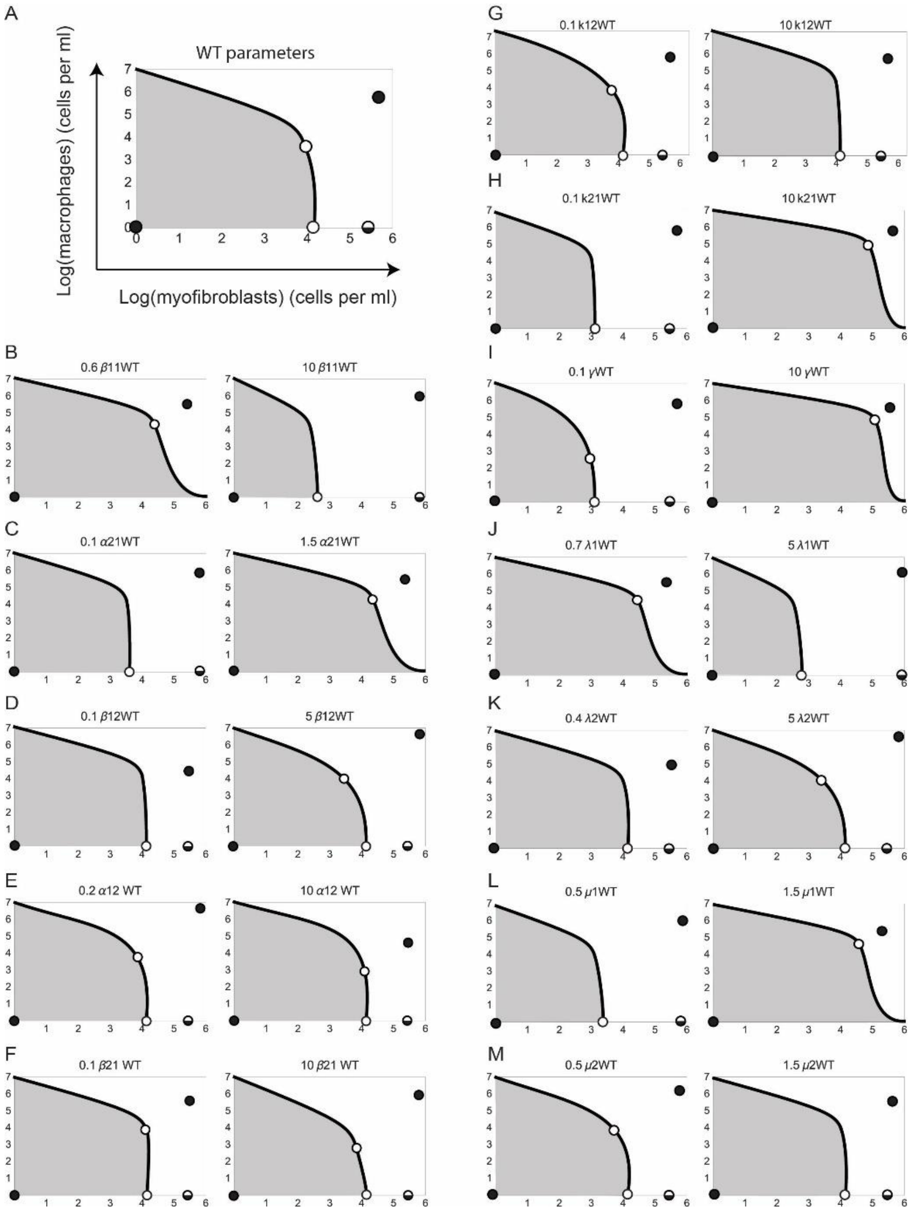


**Figure S2: The healing and hot fibrosis states are stable for a wide range of model parameters.** (A) Phase portrait with WT parameters. Stable fixed points (black dots) and unstable fixed points (white dots) are shown (the semi-stable cold fibrosis state is split black and white), as is the separatrix (black line) that marks the boundary between the basin of attraction of the healing state (gray region) and fibrosis state. (B-M) Phase portrait with one of the model parameters smaller (left panel) or larger (right panel) than the WT model parameter value. When parameters are shifted by values other than 0.1-fold and 10-fold, the shift indicates the boundary of the stable parameter range.

**The cold-fibrosis state can be made stable with altered model parameters or additional interactions.**

The cold fibrosis state is a semi-stable fixed point with the parameters that are used in Figure 1E because a small influx of macrophages leads to the hot fibrosis state. The cold fibrosis state can be made fully stable by using parameters in which the hot fibrosis state loses stability, leaving only the healing and cold fibrosis states as stable solutions (Fig S3A).

Another scenario that stabilizes the cold fibrosis state is considering a slightly modified model in which CSF production in myofibroblasts is downregulated by PDGF. In this case, perturbing cold fibrosis by adding a small amount of macrophages leads to a return to cold fibrosis and a loss of the macrophages (Fig S3B); addition of a large amount of macrophages causes flow to the hot fibrosis state (Fig S3B).

We model CSF downregulation by PDGF using the following equation for CSF:

(S3) $C\dot{S}F = \beta_1 \frac{k_1}{k_1+PDGF} mF - \alpha_1 M \frac{CSF}{k_2+CSF} - \gamma CSF$

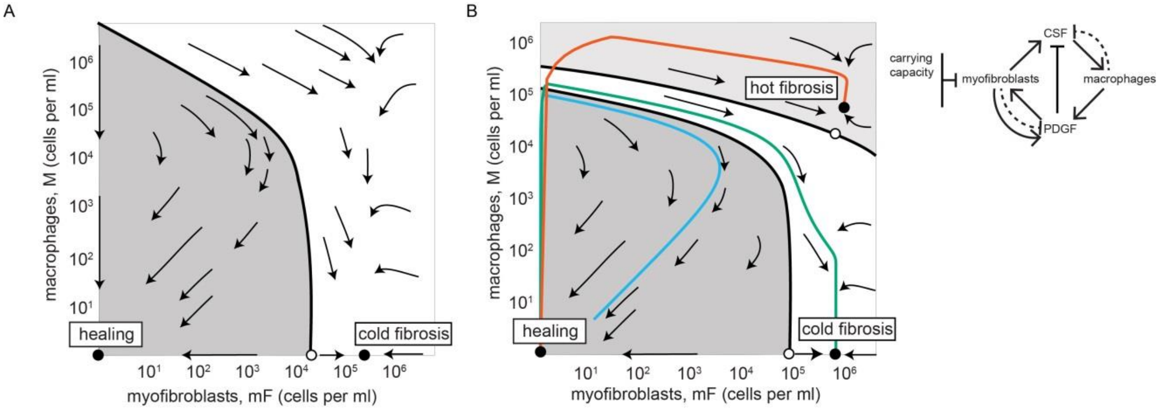


**Figure S3: The cold-fibrosis state can be made stable with altered model parameters or additional interactions.** (A) Myofibroblast-macrophage phase portrait with CSF secretion rate 100-fold lower than in Figure 1E stabilizes the cold-fibrosis state. There is no hot fibrosis state. (B) Myofibroblast-macrophage phase portrait in a circuit in which PDGF downregulates CSF expression in myofibroblast shows all three stable states. Note there are two separatrix curves, dividing the phase portrait into three basins of attraction. The middle basin (white region) flows to the cold-fibrosis state. A transient injury leads to the healing state (light blue line), an injury with intermediate duration leads to the cold fibrosis state (green line), and a prolonged injury leads to the hot fibrosis state (red line). We used the parameter values: $\alpha_1 = 0.2 \frac{molecules}{cell\ min}, \alpha_2 = 30 \frac{molecules}{cell\ min}, \beta_3 = 8 \frac{molecules}{cell\ min}, \beta_1 = 2 \frac{molecules}{cell\ min}, \lambda_1 = \lambda_2 = 2 \frac{1}{day}$. We used the values listed in Table 1 for the remaining model parameters.

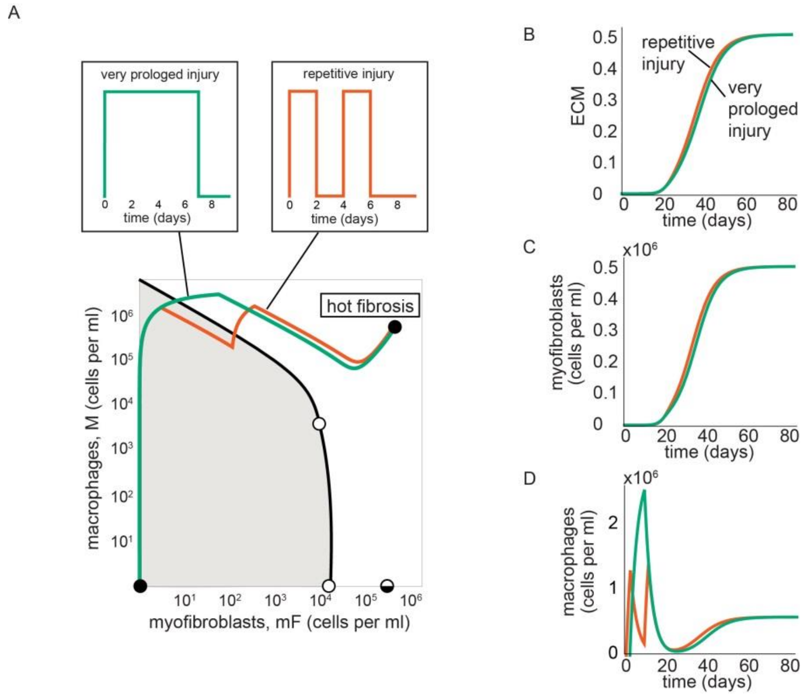


**Figure S4: Repetitive injury results in a similar response as a very prolonged injury.** (A) A 7-day inflammatory pulse and two successive 2- day pulses lead to a trajectory to the hot-fibrosis state with persistent mF and M populations (in green and red, respectively). The dynamics of ECM (B), myofibroblasts (C), and macrophages (D) following these injuries are similar especially at late times.
